## Supplementary figures and images for "Expanding the Drosophila toolkit for dual control of gene expression"

### Figure 5 source data 1

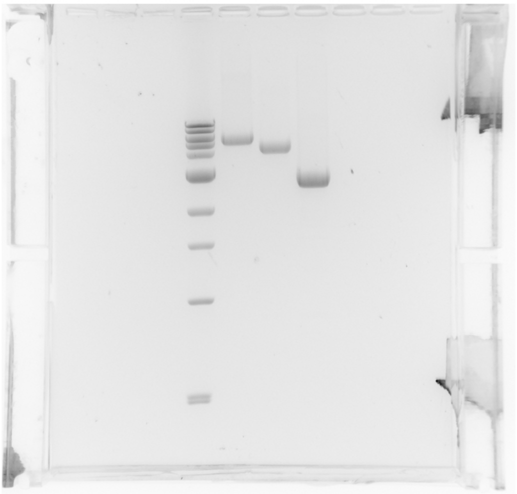

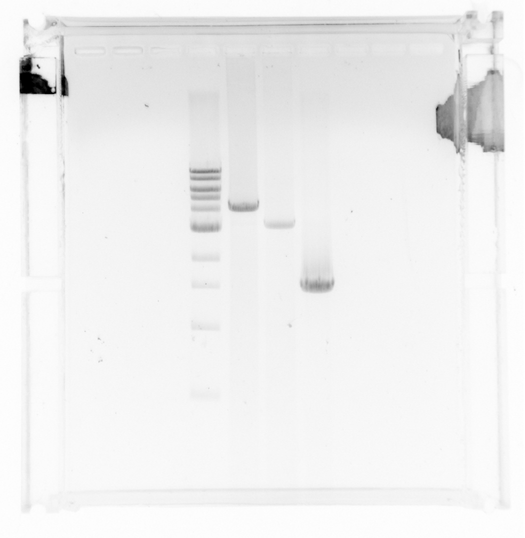

### Figure 5 source data 2

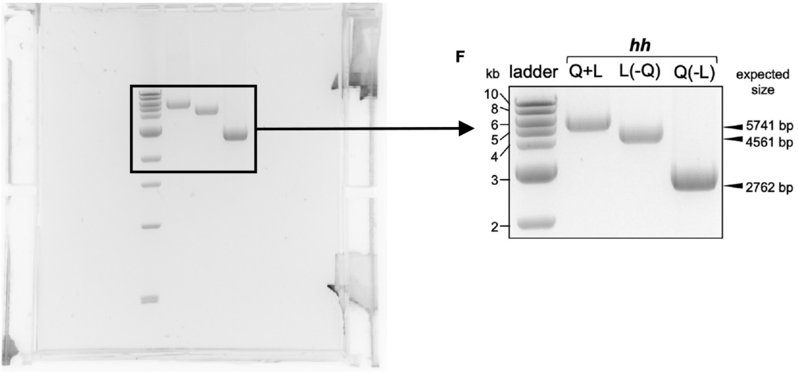

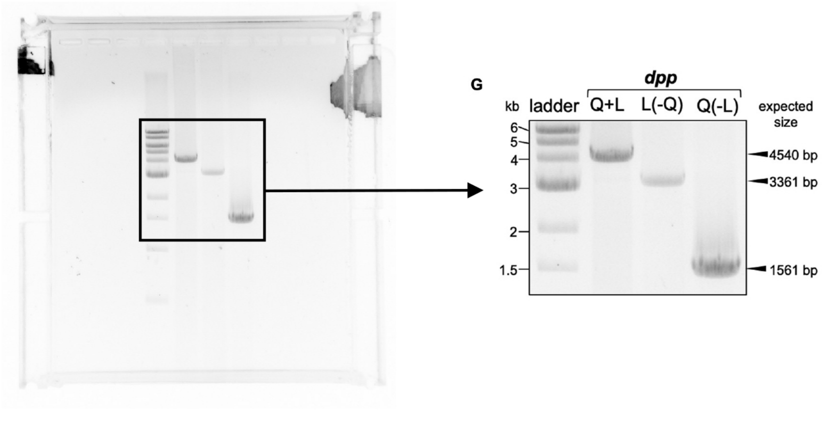
